# Thermoelectric Heat Exchange and Growth Regulation in a Continuous Yeast Culture

**DOI:** 10.1101/258962

**Authors:** Adriel Latorre-Pérez, Cristina Vilanova, José J Alcaina, Manuel Porcar

## Abstract

We have designed a thermoelectric heat exchanger (TEHE) for microbial fermentations, able to control the temperature of a microbial continuous culture, and produce electric power. The system proved able to stably maintain both the temperature and the optical density of the culture during the exponential, highly productive phase.

## 1. Introduction

A range of parameters such as temperature, pH, or substrate concentration need to be stable in order to sustain a suitable microbial growth and/or the stable biosynthesis of a bioproduct [1]. Temperature strongly affects a range of fundamental cellular processes [2, 3]; and thus keeping a microbial culture in a suitable range of temperatures is of high importance in terms of strain performance [4]. Large-scale growth of most microorganisms is accompanied by the production of heat [5], which, when large culture volumes are set, often results in an undesirable increase in the temperature of the batch culture that has to be alleviated through refrigeration [6, 7].

In a previous work, we described the first Microbial Thermoelectric Cell (MTC), a system designed for batch cultures allowing the partial conversion of microbial metabolic heat into electricity [8]. A range of industrial fermentations are carried out in continuous culture, where stable cellular densities can be maintained during long periods thanks to the supply of fresh medium, which is introduced at a rate that is equal to the volume of product that is removed from the fermenter. In this work, we aimed at the design, construction and characterization of a continuous culture system where temperature is automatically controlled and electric power is constantly obtained during all the fermentation process. To do that, we envisaged, constructed, and set in place a ThermoElectric Heat Exchanger (hereafter called TEHE), a device based on the Seebeck effect, which allows a fine control of temperature and fresh medium input and thus microbial growth-while electric power is produced.

## 2. Materials and Methods

### 2.1. Experimental set-up

A medium-scale continuous culture of budding yeast Saccharomyces cerevisiae strain D170 (kindly provided by Prof. Emilia Matallana, IATA, Valencia, Spain) in YPD medium supplemented with 18 % sucrose was set up in the laboratory as schematically represented in Fig. 1A. The TEHE consisted of two aluminum pipes of squared section and a serial connection of ten thermogenerators (MCPE-071-10-13, Multicomp) placed in direct contact with the pipes. The whole device was thermally insulated with expanded polystyrene (EPS) and polyurethane foam spray (Silicex Fischer, Fisher Ibrica, Tarragona, Spain) (Fig. 1B). The TEHE was coupled to a thermally isolated 40 L Dewar flask (Scharlab, Barcelona, Spain) combined with a MM-1000 overhead anchor stirrer (Labnet International, Edison, NJ, USA), and two peristaltic pumps (Lambda Laboratory Instruments, Baar, Switzerland), which were programmed to control the flow of fresh and wasted medium.

**Figure 1:**
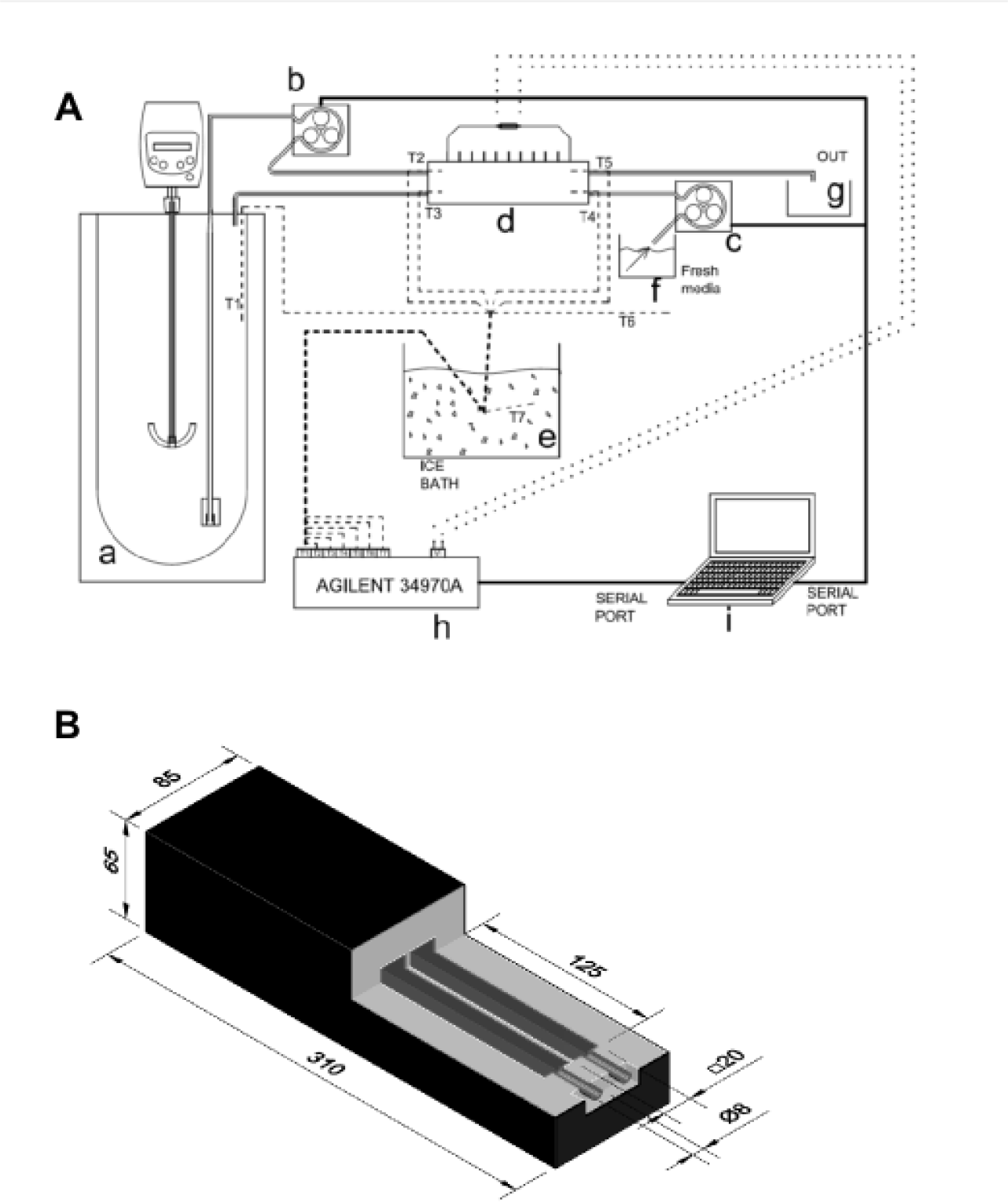
**(A)** Schematic representation of the continuous culture system. (a) Thermally insulated fermenter; (b, c) peristaltic pumps; (d) ThermoElectric Heat Exchanger (TEHE); (e) ice bath for the compensation of temperature measurements; (f) refrigerated fresh medium tank; (g) wasted medium tank; (h) data logger; and (i) PC with a software (Rodrguez-Barreiro et al., 2013) for data recording and automatic control of the peristaltic pumps. All temperature measurements were performed with T-type thermocouples. **(B)** Tridimensional representation of the TEHE constructed in this work. Sizes given in mm.

### 2.2. Data Acquisition, Monitoring and Recording

The whole system was connected to a PC in order to record temperature values as well as output electrical current. Temperature measurements were performed by thin T-type thermocouples inserted into the different parts of the system and connected to a PC through a data logger. As in previous studies [8], the connections between the thermocouples and the data logger were performed on an ice-water mixture to take into account the unwanted background electric voltage, due to the junction of dissimilar metals in the thermocouple-data logger connection. Temperature and electrical current records were taken every 6 minutes throughout the experiment. Both feed and effluent flows were automatically modulated during all the experiment with the LabVIEW control software.

### 2.3. Mathematical modelling

The global thermal resistance (*Rg*) and the whole heat capacity (*mc_P_*) of the fermenter were estimated from Equation 1:

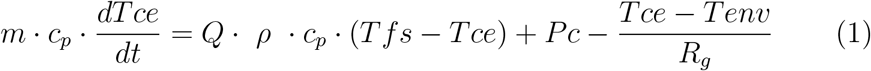

Where *m, c_P_*, and *Tce* are broth mass, specific heat, density, and temperature, respectively, whereas *Q* is the flow rate and *Pc* is the metabolic heat produced by yeasts. *Tfs* and *Tenv* correspond to the fresh medium entering the fermenter and room temperatures, respectively. Equations 2 and 3, describing a Logarithmic Mean Temperature Difference (*LMTD*) model of a heat exchanger [9], were used to mathematically characterize the TEHE:

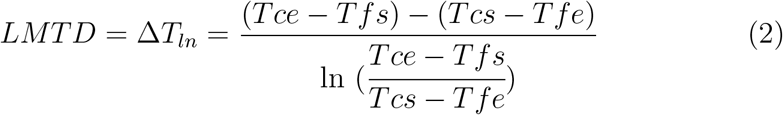

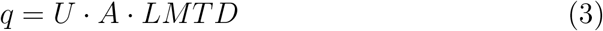

Being *Tce* and *Tfe* the temperature of the hot and cold inlet flows, respectively; and *Tcs* and *Tfs*, the temperature of the hot and cold outlet flows, respectively. The heat flow (*q*) between the hot and the cold pipe depends on the temperature of the fluids entering the exchanger, the global heat transmission coefficient (*U*), and the heat exchange surface (*A*).

In order to obtain an estimation of the *UA* constant, *q* was first calculated from Equation 4 in an experiment where two water flows at known temperatures were entered in the TEHE.

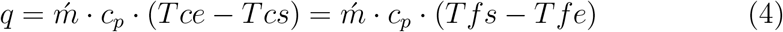

Where ḿ is the inlet mass flow rate (mass of water entering the TEHE per unit of time), and c_p_ is the specific heat of water.

## 3. Results and Discussion

The output of a typical experiment carried out in the continuous culture system set as described above is shown in Fig. 2. The broth (35 L) was inoculated with 700 mL (1:50) of an overnight yeast culture, and cultivated in the thermally isolated flask under shaking (180 rpm). Broth temperature rose in an exponential fashion and reached 35 °C after 24 h, (Fig. 2A). At this point, culture temperature was kept constant by means of introducing fresh, cool medium in the fermenter (feed flow) at the same rate that wasted (warm) medium was extracted (effluent flow), in such a way that the volume of the culture did not change during the experiment. The heat flow through both aluminum pipes warmed the fresh medium (from 18 up to 22°C, approximately) prior to its entrance to the fermenter (Fig. 2B); and, reciprocally, cooled the waste warm (temperature at the TEHE input, around 30°C) down to around 25 °C, approximately. As a result, a rather stable voltage of 1-1,3 V was recorded (Fig. 2C). In order for the TEHE to produce the maximum electric power, a load resistance of 120 Ω was coupled to the terminals of the thermogenerators, yielding 10-12 mW. When the feed and effluent flows were halted and the TEHE was not used, the temperature of the broth started rising immediately, peaked at 42 °C, and then started to drop (Fig. 2A).

**Figure 2:**
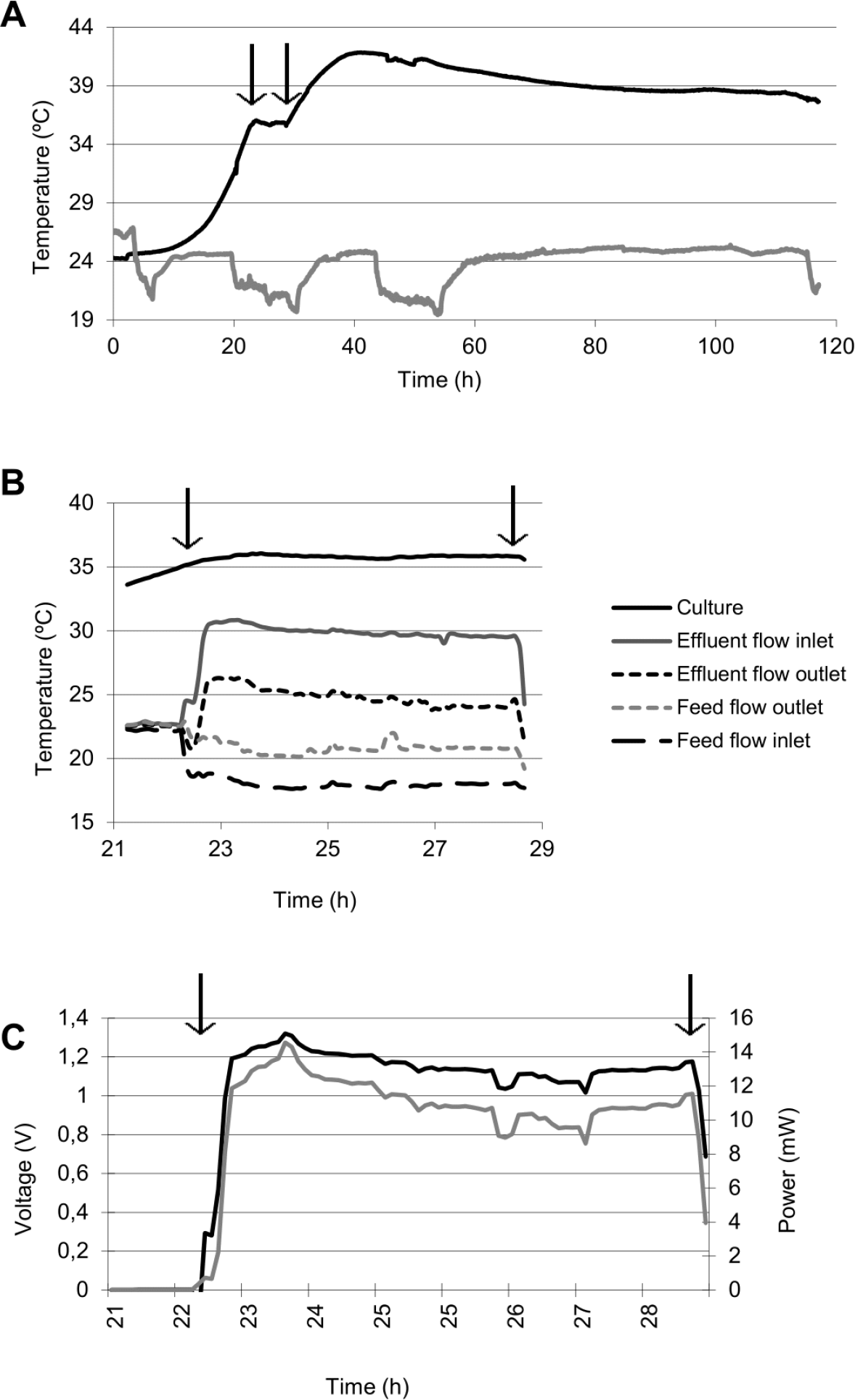
**(A)** Evolution of broth (black line) and room temperatures (grey line) in a typical experiment. Arrows indicate the period of time when the TEHE was connected. **(B)** Changes in the temperature of inlet and outlet flows of the TEHE. **(C)** Voltage (black line) and power (grey line) production in the TEHE.

The thermal behavior of the two main components of the system (the fermenter and the TEHE) was experimentally characterized and mathematically modeled. Following a simplified experimental set up where no fresh medium (*Q*=0) nor cells (*Pc*=0) were introduced in the fermenter, an identification assay was performed to estimate *Rg* and *mc_p_*, obtaining values of 5,92 K/W and 146,547 kJ/K, respectively. The UA constant was estimated also estimated as explained in the Materials and Methods section, yielding a value of 1,039 W/K.

The evolution of the yeast culture was studied in a typical experiment where optical density (OD) at 600 nm was periodically measured. As shown in Fig. 3, yeast population and temperature during the first part of the experiment exhibited a similar pattern. When the TEHE was connected and temperature was kept constant, the OD_600_ of the broth was relatively stable at around 8, indicating that the number of cells present in the fermenter was maintained stable despite the large flow of broth removal (2,4 L/h on average).

**Figure 3:**
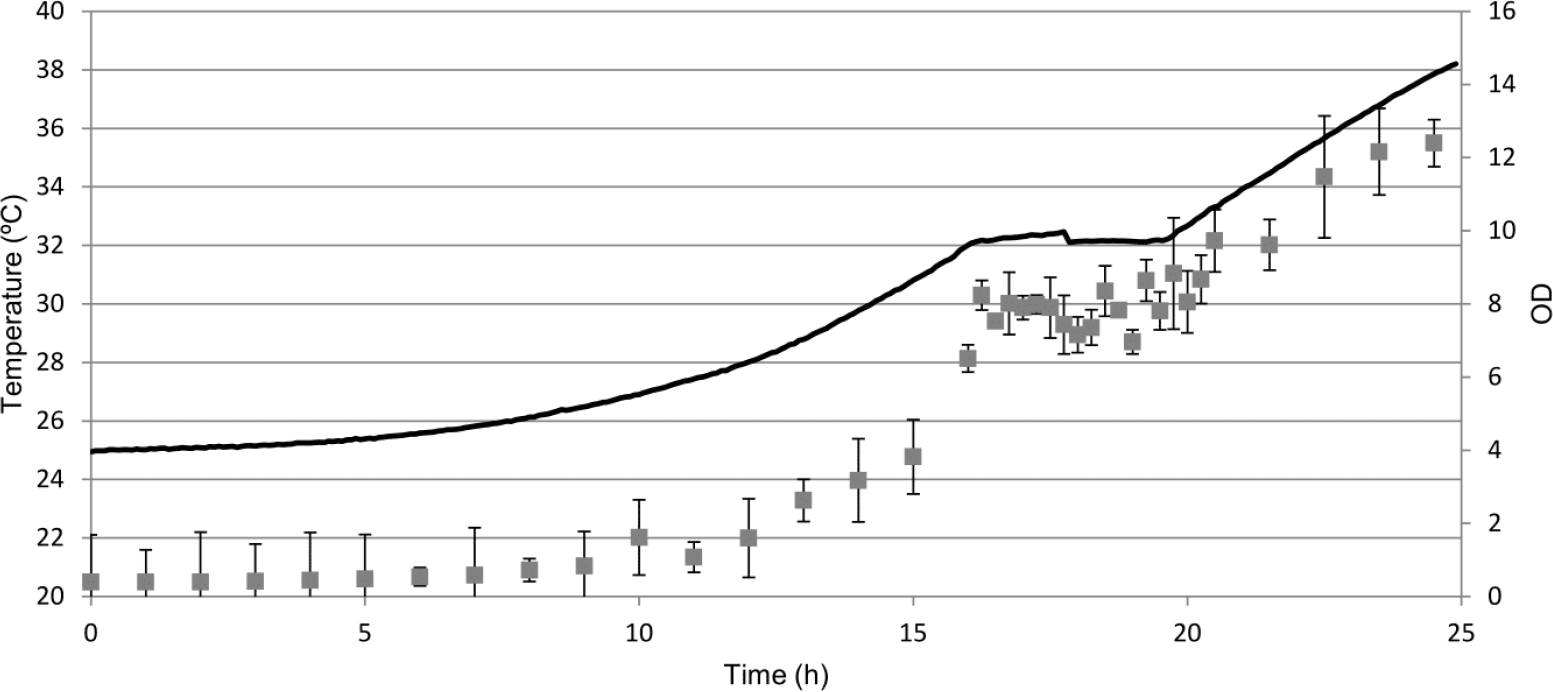
Evolution of broth temperature (black line) and optical density at 600 nm (grey squares). Arrows indicate the interval of time when the TEHE was connected. Error bars show the standard deviation of three independent measurements.

Taken together, our results prove the ability of this thermoelectric heat exchanger-based system to autonomously regulate the broth temperature and to produce electric power by harvesting metabolic heat.

In our prototype, autonomous heating of the culture was achieved and reached values (42 °C) well beyond optimal temperatures for budding yeast. Lower, industrially friendly temperatures could be constantly maintained by means of an automatic equilibrium between the flow of fresh and product-containing media through the TEHE. This resulted in the production of a significant electric power during all the process. In addition, biomass concentration proved to be constant when the temperature was controlled. This is of key importance, since industrial bioprocesses require stable temperatures in order to maintain a constant output of a given product [10].

The generation of electric power in heat exchangers through the Peltier-Seebeck effect has been previously reported [11-13]. However, this is, to the best of our knowledge, the first time that a heat exchanger with thermogenerator ability has been coupled to a bioprocess. The difference of temperatures that can be achieved between both sides of the thermogenerators is obviously limited by the narrow range of temperatures mesophiles can tolerate; and, therefore, the electric power that can be produced is lower than that of other type of industrial processes [14]. Albeit low, TEHE-based power production proved twice more efficient compared to our previous MTC designed for batch culture. Indeed, a 40-fold increase in electrical power production was obtained in the TEHE compared to MTC, which was 20-fold smaller [8].

Our results are the first step towards controlling microbial growth towards thermoelectric exchangers producing electric power. Scaling up fermentation volumes would accordingly increase electrical power production, which might allow a at least-partial self-control of the culture based on its temperature, with the TEHE contributing to power the peristaltic pumps that regulate broth renovation rates.

## Acknowledgments

We are very grateful to Julin Heredero, from ICMUV (Institut Cincia Materials Universitat Valncia) for manufacturing the TEHE pipes. This work was funded by the Valoritza i Transfereix program (CPI-13-128) from the University of Valencia. The technology described in this work is subjected to industrial protection (Patent Cooperation Treaty reference number: PCT/ES2013/000212). The authors have prepared the patent and the registration in collaboration with the Research Transfer Office (OTRI) of the University of Valencia (contact person, Marta Garcs).

